# Sub-kb resolution Hi-C in D. *melanogaster* reveals conserved characteristics of TADs between insect and mammalian cells

**DOI:** 10.1101/164467

**Authors:** Qi Wang, Qiu Sun, Daniel M. Czajkowsky, Zhifeng Shao

**Affiliations:** Shanghai Center for Systems Biomedicine, Shanghai Jiao Tong University, Shanghai 200240, China; State Key Laboratory for Oncogenes and Bio-ID Center, School of Biomedical Engineering, Shanghai Jiao Tong University, Shanghai 200240, China

**Author notes:** Q.W. and Q.S. contributed equally to this work.

## Abstract

Topologically associating domains (TADs) are fundamental elements of the 3D structure of the eukaryotic genome. However, while the structural importance of the insulator protein CTCF together with cohesin at TAD borders in mammalian cells is well established, the absence of such co-localization at most TAD borders in recent Hi-C studies of *D. melanogaster* is enigmatic, raising the possibility that these TAD border elements are not generally conserved among metazoans. Using *in situ* Hi-C with sub-kb resolution, we show that the genome of *D. melanogaster* is almost completely partitioned into more than 4,000 TADs (median size, 13 kb), nearly 7-fold more than previously identified. The overwhelming majority of these TADs are demarcated by pairs of *Drosophila* specific insulator proteins, BEAF-32/CP190 or BEAF-32/Chromator, indicating that these proteins may play an analogous role in *Drosophila* as that of the CTCF/cohesin pair in mammals. Moreover, we find that previously identified TADs enriched for inactive chromatin are predominantly assembled from the higher-level interactions between smaller TADs. In contrast, the contiguous small TADs in regions previously thought to be unstructured “inter-TADs” are organized in an open configuration with far fewer TAD-TAD interactions. Such structures can also be identified in some “inter-TAD” regions of the mammalian genome, suggesting that larger assemblages of small self-associating TADs separated by a “burst” of contiguous small, weakly associating TADs may be a conserved, basic characteristic of the higher order folding of the metazoan genome.

## INTRODUCTION

It is now widely recognized that the 3D structure of the genome plays a fundamental role in many nuclear processes, from cellular differentiation to transcriptional regulation to DNA replication and repair (1-6). Methods derived from chromosome conformation capture (3C) (7), such as 5C (8) and Hi-C (9), have proven particularly instrumental in this regard, revealing topologically associating domains (TADs) within which genomic loci are found to contact each other more frequently than those between adjacent TADs or in adjacent de-condensed, unstructured “inter-TAD” regions between TADs. Such TADs and inter-TAD regions have now been observed in most eukaryotic cells, suggesting that these are basic structural elements of the genomic architecture (10, 11).

In mammalian cells, there are now a number of studies demonstrating that the insulator protein CTCF and cohesin co-localize to the borders of many TADs (12-14), with CTCF directly interacting with specific DNA sequences and cohesin mediating long-range chromosomal interactions (13). Based on studies of targeted deletion of specific CTCF binding sites, the presence of CTCF and cohesin at TAD borders has been shown to be pivotal for the formation of TADs (14-17). However, whether these proteins or their homologues play a similar function in other metazoan cells is not clear. In particular, for the model organism *Drosophila melanogaster*, recent Hi-C studies failed to demonstrate a significant enrichment of either dCTCF (the CTCF homologue in *D. melanogaster*) or cohesin at TAD borders (18, 19). Such a discrepancy is perplexing as one might expect such functionally important complexes to be conserved given the overall conservation of many basic biological and physiological processes between mammals and *D. melanogaster* (20).

Here we re-investigated the global structure of the *D. melanogaster* genome using *in situ* Hi-C at high depth to achieve a restriction-site limited “map resolution” of ∼200 bp. At this higher resolution, we find that there are many more (nearly 7-fold) TADs resolvable in this genomic structure than previously identified. More importantly, there is a strikingly high enrichment of pairs of the insulator proteins, BEAF-32/CP190 or BEAF/Chromator, at the TAD borders, analogous to the enrichment of CTCF/cohesin pairs at TAD borders in mammalian cells. Further, while most of the previously identified TADs, primarily enriched for inactive chromatin, are now resolved as higher-order assemblages of smaller TADs, unexpectedly, the previously identified inter-TAD regions, thought to be unstructured, are actually composed of a string of well-defined small TADs with limited near-range inter-TAD contacts, a feature that can also be identified in mammalian cells. Taken together, these results strongly suggest that several of the most basic features of the higher order genome architecture are conserved from insects to mammals.

## Results

### The *D. melanogaster* genome is fully partitioned into contiguous TADs

Since the chromosome structure in eukaryotic cells is known to change significantly during the cell cycle (21), we sought to minimize the variability in our examination of the genomic structure of the model eukaryote, *D. melanogaster*, by studying cells that were arrested at the G1/S boundary. To this end, we incubated S2R+ cells (22), a well-studied cell line derived from the late embryo, with hydroxyurea, which is an effective inhibitor of eukaryotic DNA replication (23) (Fig. S1A). We performed *in situ* Hi-C using the 4-cutter restriction enzyme, DpnII, following an established protocol (24) with minor changes (Materials and Methods, Supplementary Materials). The median length of the DpnII restriction fragments in this genome is 194 bp. Sequencing the Hi-C library generated 695 million raw reads, which yielded 255 million high-quality read-pairs after all filtration steps (Supplementary Materials). To evaluate the reliability of this data, we also performed *in situ* Hi-C on a biological duplicate, sequencing to a lower depth of 253 million raw reads that yielded 98 million valid pairs. The two datasets were highly correlated (Pearson’s correlation, *r* = 0.98) (Supplementary Materials). Consequently, for all further analysis, we combined both datasets to finally obtain 353 million pair-end reads with a maximal estimated “map resolution” of ∼200 bp, as calculated following Rao et al (24).

To ensure the validity of our data, we generated a contact map at a lower resolution (20 kb) and compared it with that obtained previously from the highly related, S2 cells that was of this resolution (19). Using the Armatus software to annotate TADs (19, 25), we identified 612 TADs that exhibited a median size of 140 kb, bordered by inter-TAD regions of a median size of 40 kb (Table S1). These results are in excellent agreement with this earlier study (19). In fact, the precise location of our TAD borders exhibited a high degree of overlap (81.3%) with those identified in the previous work (Supplementary Materials, Fig. S2A and B). We also confirmed the lack of significant co-localization of dCTCF or cohesin at these TAD borders (Supplementary Materials, Fig. S2C). Thus, at this lower resolution, our data and analysis agree substantially with this earlier study.

However, when our data are examined at the higher, restriction fragment-limited resolution, it is immediately apparent that there are in fact many small TADs within both previously defined TADs and, notably, within the so-called “inter-TAD” regions (Fig. 1A). To avoid confusion, we will henceforth refer to those TADs identified at 20 kb resolution as “super-TADs” as they are in general much larger than those observed at the fragment-limited resolution, which we will refer to as “TADs”. Likewise, we will refer to the regions between the super-TADs as “inter-super-TADs”.

**Figure 1.**
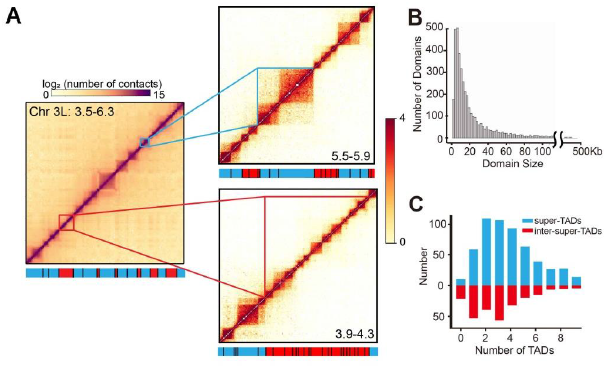
The Drosophila genome is fully partitioned into contiguous TADs including within previously annotated “inter-TADs” regions. (A) Heatmaps from the left arm of chromosome 3. The left panel shows a heatmap of a 2.8 Mb region of this arm at 20 kb resolution, revealing well-defined super-TADs (blue bars at the bottom) and inter-super-TADs (red bars at the bottom), consistent with previous findings (19). However, at the highest resolution permitted by the data, the heatmap shows that both the super-TADs (right upper panel) and the inter-super-TADs (right lower panel) are composed of small contiguous TADs. The blue (red) bars in these panels now refer to TADs (inter-TADs). (B) The size distribution of the TADs annotated from the fragment-limited resolution map. The median size of the TADs is 13 kb, much smaller than the size (140 kb) identified from previous lower resolution data. (C) The number of TADs within the super-TADs (blue bars) and inter-super-TADs (red bars). The super-TADs are found to consist of a range of TADs, mostly between 2 to 4, while there are generally between 1 to 4 TADs within the inter-super-TADs, where previous work concluded that there were no TADs.

In total, we identified 4,123 TADs that range in size from 3 kb to 460 kb, with a median size of 13 kb, that altogether cover almost the entire (>92%) 130 Mb non-repetitive *D. melanogaster* genome (Fig. 1A and B, Fig. S3). As shown in Fig 1, the super-TADs are now found to be subdivided into most frequently 2 to 4 small TADs (median size, 16 kb, Fig. 1C). By contrast, the inter-super-TADs that were previously considered largely devoid of identifiable organization are shown to completely consist of generally 1 to 4, slightly smaller TADs (median size, 9 kb, Fig. 1C).

A striking feature of the distribution of these TADs, whether associated with a super-TAD or inter-super-TAD, is that most (75.4%) of the borders between adjacent TADs localize to the same restriction fragment (Fig. S3, Table S2). That is, at the resolution limited by the size of the restriction fragments, the TADs are essentially contiguous, without an unstructured region in between, illustrating that, unlike what was concluded from lower resolution maps, there are essentially no extended stretches of “inter-TADs” across the genome.

### Demarcation of TADs by specific pairs of insulator proteins

Yet, even with these more precisely defined borders, a comparison with the known locations of dCTCF or cohesin subunits showed an absence of significant enrichment at TAD borders (Fig. S4C), consistent with previous work showing that neither protein defines border elements in *D. melanogaster* (18, 19). However, since this organism contains many other insulator proteins (26, 27), we reasoned that other insulator proteins might function as analogues of CTCF/cohesin in this organism instead. An early study identified two classes of insulator proteins in *Drosophila* embryos (28): Class I (that includes BEAF-32 and CP190) and Class II (that includes only Su(Hw)). Surprisingly, using the binding site locations defined in this earlier work, we found an exceptionally high co-localization of Class I insulator proteins at these narrowly defined TAD borders (Fig. 2A, Fig. S4B). By contrast, the Class II insulator protein was not significantly associated with TAD borders (Fig. S4A and B).

**Figure 2.**
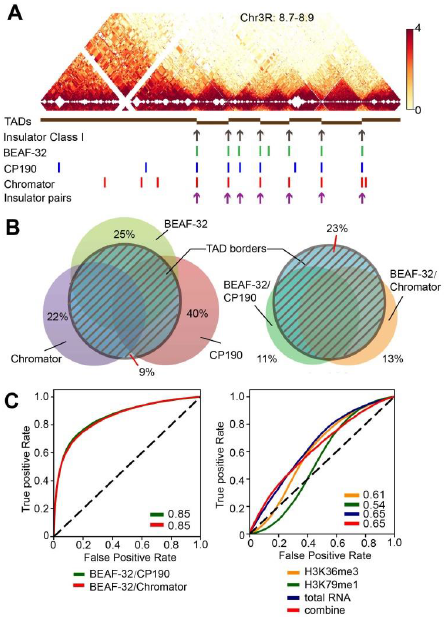
The TADs are demarcated by pairs of insulator proteins. (A) The locations of known Drosophila insulator proteins, together with the TADs identified in this work, are shown for a 200Kb segment of chr3R. The positions of Class I insulator proteins (that includes BEAF-32 and CP-190), as obtained from Flybase, are highly localized to the TAD borders. Also shown are the peak locations of the individual insulator proteins (BEAF-32, CP-190, and Chromator) characterized in the modENCODE project that are enriched at the TAD borders. However, only pairs of BEAF-32/CP190 or BEAF-32/Chromator are found to be exclusively associated with the TAD borders. (B) Venn diagram showing the genome-wide co-localization of these insulator proteins and insulator protein pairs at the TAD borders. There is a significantly greater exclusive association of the insulator protein pairs at TAD borders than of the individual proteins separately. (C) The enrichment of the insulator protein pairs at the TAD borders is further validated by the examination of the extent to which the locations of the insulator protein pairs are predictive of TAD borders using logistic regression models. Shown are the receiver operating characteristics curves (ROC), with the area under the curve (AUC), reflecting the predictive power indicated in the inset of each panel. Both pairs of insulator proteins are highly predictive of TAD borders (left panel), while transcriptionally active epigenetic modifications or transcriptional levels (right panel) are poorly predictive.

To determine the enrichment of individual insulator proteins, we analyzed the location of all insulator proteins profiled in the modENCODE project in S2 cells (namely, dCTCF, BEAF-32, Su(Hw), GAF, ZW5, CP190, Chromator and mod(mdg4)). We found that BEAF-32, CP190 and Chromator are each significantly enriched at the boundaries of the TADs (Fig. S4C) while no other insulator protein exhibits such significant enrichment at these TAD borders (Fig. S4C). Overall, more than 91% of all TAD borders contain at least one of these three proteins (Fig. 2B), an enrichment that far exceeds what would be expected from a random distribution (Fisher’s exact test, p value < 2.2e-16).

However, we also found that, as with CTCF and cohesin in mammalian cells (13,14, 29), each of these insulator proteins is found at many other locations in addition to TAD borders (Fig. 2B). Since previous work has shown that BEAF-32 binds to specific DNA sequences (26, 30) and CP190 and Chromator both bind to BEAF-32 and mediate long-range chromosomal contacts (30), we examined if there is a greater degree of exclusivity at the TAD borders of pairs of insulator proteins (BEAF-32/CP190 or BEAF-32/Chromator) than what is observed with the individual proteins. Indeed, we found that 74% of pairs BEAF-32/CP190 or BEAF-32/Chromator localize to the TAD borders, and conversely, 77% of the borders localize to the binding sites of these pairs (Fig. 2B). This striking correlation holds true over a wide range of the Armatus TAD annotation parameters (Fig. S4D).

We further validated this enrichment by examining the extent to which the positions of these protein pairs alone could predict the location of TAD borders using logistic regression, as described in previous work (19). We found that regression based on the locations of the pairs of BEAF-32/CP190 or BEAF-32/Chromator is highly predictive of a TAD border (Fig. 2C). By contrast, similar analysis with active transcription markers (H3k26me3 and H3k79me1) or total RNA or their combination, which have been previously suggested to be generally associated with TAD boundaries in *Drosophila* (19), are substantially less predictive of TAD borders (Fig. 2C). Thus, BEAF-32/CP190 and BEAF-32/Chromator may be defined as bona fide TAD border elements in *D. melanogaster.*

### Chromatin state correlates with higher-order interactions between TADs to form super-TADs

Previous work suggested that histone modifications are a major driving factor for TAD formation in *Drosophila* and other eukaryotes (19, 31). To examine the relationship between chromatin state and the TADs identified here, we first classified the TADs according to the enrichment of 15 histone modifications and non-histone chromosomal proteins within each TAD using k-means clustering (32), identifying eight different types that could be broadly grouped into four major types of TADs: those enriched with active, inactive, or polycomb-associated chromatin marks/proteins, and those without any of these features (“undetermined”) (Fig. 3A, Fig. S6A). Consistent with previous work, we found that 83% of TADs enriched for inactive chromatin localize within super-TADs, while 81% of TADs enriched for active chromatin localize within inter-super-TADs, a highly non-random distribution (Fisher’s exact test with *p* value < 2.2e-16). We note, though, that such a correlation is not present at the TAD level.

**Figure 3.**
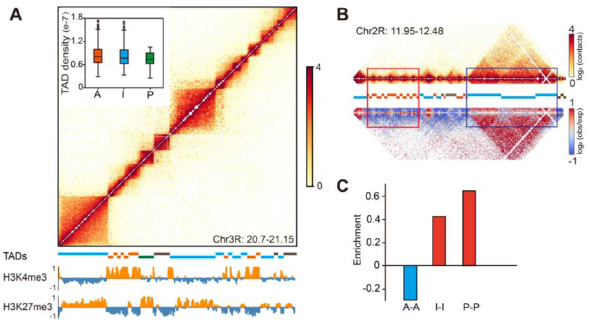
Epigenetic modifications only correlate with higher order folding of the TADs but not the folding of individual TADs. (A) The TADs could be classified into four major types according to the enrichment of 15 histone modifications and non-histone chromosomal proteins within each TAD (Supplementary Materials). Shown is an example of the distribution of these types with active (orange bar below the heatmap), inactive (blue bar), polycomb (green bar), and undetermined (grey bar) chromatin within the TADs in a 350 kb region of chr2L. Inset: The extent of DNA condensation within the TADs, as determined from the sum of contact frequencies between loci within the TAD, is the same, regardless of the type of chromatin that is enriched within the TAD (A, active; I, inactive; P, polycomb). (B) Active and inactive TADs exhibit dramatically different tendencies to interact with neighboring TADs of the same type. The upper heatmap shows the positions of the TADs, while the lower heatmap shows the significance of the observed contacts, with those colored red (blue) exhibiting much greater (lower) interaction strength than expected by chance (materials and methods) in a 530 kb region of chr2R. (C) The relative interaction strength (as shown in (B)) between pairs of TADs, indicating that TADs containing active chromatin generally tend to avoid interacting with each other while those containing inactive or polycomb chromatin frequently interact with each other.

However, we found that, at least over the distances over which a comparison can be made (Materials and Methods), the TADs enriched for inactive chromatin exhibit the same level of DNA condensation as those enriched for active chromatin, as determined from the sum of contact frequencies within the TADs, as previously (19) (Fig. 3A, inset). By contrast, there is a significant enrichment of inactive-inactive or polycomb-polycomb inter-TAD contacts between neighboring TADs and a strong depletion of active-active TAD contacts over what would be observed by chance (Fig. 3B and C). Further, overall, the TADs within the super-TADs make far more frequent contacts with other TADs within the super-TAD while those within inter-super-TADs contact each other make significantly fewer contacts with neighboring TADs (Fig. S5). Thus, overall, this analysis suggests that the chromatin state is a driving force not for condensation within TADs, but rather for interactions between the TADs responsible for the folding into the higher-order super-TAD structures.

### Comparison between high resolution Hi-C maps reveals conserved features between *D. melanogaster* and mammals

An unmistakable feature of the genomic structure revealed by this high resolution map is that essentially all of the genome is folded into TADs, with more highly-ordered super-TADs separated by open regions of smaller TADs. To determine whether these structural details are only characteristics of the fly genome, we sought for evidence of these features in previously studied mammalian cells. While, to our knowledge, there is no published Hi-C study of synchronized mammalian cells to the resolution in our work, we re-examined the Hi-C data from asynchronous human lymphoblastoid (GM12878) cells with 1 kb resolution (24). As shown in Fig. 4, there were indeed many small (median size, 30 kb) contiguous TADs readily identifiable within previously defined inter-TADs. Some of the borders of these smaller TADs are also binding sites of CTCF and cohesin (Fig. 4). This earlier work noted several occurrences of larger TADs that, like the super-TADs found in *Drosophila*, are composed of smaller TADs (24). Thus, the general TAD-level organization observed in *D. melanogaster* may also be a conserved feature of the genomic structure of mammals as well.

**Figure 4.**
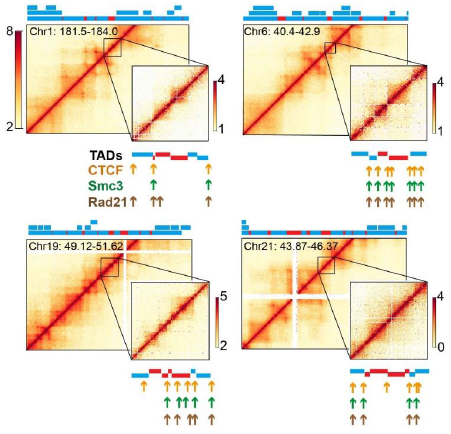
The human genome is also partitioned into contiguous small TADs within previously described “inter-TAD” regions at least in part. Shown are four examples of Hi-C data of GM12878 lymphoblastoid cells determined by Rao et al. (24). In each panel, the main figure is the heatmap of the indicated chromosomal position at 5 kb resolution, with the domains annotated by these authors indicated by the color bars above each figure. Note that there were smaller TADs within larger TADs identified in this previous work, reflected in the three different levels in the annotated TADs. The bars colored red reflect the inter-TAD regions while those colored blue are the TADs. The inset of each panel is an expanded region of an inter-TAD region showing that they consist of many smaller TADs, similar to what is observed in *D. melanogaster*. The TADs annotated using the Armatus software are shown below the heatmap. Also shown are the locations of CTCF (orange arrows) and cohesin components (Rad21 and Smc3, green and brown arrows respectively), as determined previously (24) showing that many of the borders of the smaller TADs identified here within these regions also co-localize with CTCF/cohesin.

## Discussion

Using *in situ* Hi-C with ∼200 bp resolution, we have examined the 3D organization of the *D. melanogaster* genome. We have found that this genome contains many more TADs than previously thought, most of which are smaller than what could be resolved in previous Hi-C studies. What emerges from an analysis of this high resolution map is that the genome structure generally consists of alternating stretches of two different types of TADs that differ slightly in size (9 kb vs 16 kb), but differ more significantly in the degree to which the TADs engage in inter-TAD interactions, with those within the super-TADs making more extensive inter-TAD contacts. These self-associating TADs within the super-TAD are highly enriched for inactive chromatin, while the weakly interacting TADs within the inter-super-TADs are predominantly enriched for active chromatin. We also performed Hi-C with asynchronous cells, finding similar results (though with less well-defined TAD borders, Supplementary Materials, Fig. S1B and C). Further, we re-analyzed data from a recently published Hi-C study of asynchronous Kc167 cells (18), and found generally similar results (Supplementary Materials, Fig. S4E). Thus, these are basic structural properties of the genome of *Drosophila* cells, regardless of cell cycle stage.

The finding that the inter-super-TAD region is completely partitioned into contiguous small TADs is highly unexpected based on previous Hi-C studies in both *Drosophila* and mammalian cells (10, 33, 34). In these, this unstructured open chromatin organization was suggested to be a consequence of active chromatin and active transcription, which was suggested to be generally inhibitive of well-organized genomic structures (10, 19). However, our findings strongly suggest that chromatin state does not play a primary role in the formation of TADs: (i) the inter-super-TAD region in fact contains many TADs (Fig. 1A); (ii) active chromatin and active transcription are not highly predictive of TAD borders (Fig. 2C); (iii) these TADs are not solely enriched for a single epigenetic type (Fig. S6A); and (iv) the TADs enriched for a specific chromatin state are found to exhibit a significant degree of heterogeneity of the modification contained within (Supplementary Materials, Fig. S6E). While the underlying mechanisms responsible for TAD formation remains to be determined and may differ between active and inactive chromatin, our results are consistent with chromatin state influencing higher-order folding of the TADs into super-TADs. Further, our findings instead suggest that components at the TAD border may play a more determinative role in TAD formation, since regardless of the chromatin type contained therein, the chromatin is similarly compacted (Fig. 3A, inset).

Our work also provides convincing evidence that the defining feature of the overwhelming majority of the TAD borders is the co-localization of pairs of specific insulator proteins, BEAF-32/CP-190 or BEAF-32/Chromator. These pairs are enriched at all TAD borders, regardless of the type of chromatin contained within the adjacent TADs (Supplementary Materials, Fig. S6D). There are many other insulator proteins in this organism (26, 27), but none show significant enrichment at these borders (Fig. S4C). Thus, the enrichment of insulator proteins at TAD borders is not a general property of all insulator proteins in this organism and thus, probably, of insulation *per se*. Conversely, we speculate that it is the presence of these pairs of insulator proteins that determines the presence of a TAD border.

The presence of a pair of proteins at the TAD border, one of which specifically binds DNA and the other that mediates long-range chromosomal interactions, which we observe here in *D. melanogaster,* is strikingly similar to what is observed at many TAD borders in mammalian cells with CTCF/cohesin. Thus, we suggest that BEAF-32/CP-190 and BEAF-32/Chromator are functional analogues of CTCF/cohesin as TAD border elements. Likewise, since CTCF and cohesin are not found at all TAD borders in the mammalian genome, there may be other similar protein pairs (35) that are also functionally analogues of CTCF/cohesin at these other TAD borders in mammalian cells. Recent work has indeed identified many other “architectural” proteins in mammals, some of which appear to be enriched in some TAD borders (36, 37).

We have also noted other structural details of TADs that may be conserved between *D. melanogaster* and mammals, most notably that the genome structure may generally consist of alternating stretches of self-interacting small TADs and weakly-interacting small TADs. The conservation of this particular pattern likely reflects necessary functional utility, perhaps providing the combination of order yet flexibility needed for a range of genomic functions (4, 38).

In conclusion, *Drosophila* is considered as a model eukaryote whose study provides direct information of basic biological processes in higher-level organisms (20, 39). Our work here extends these similarities to that of genome structure, which further underscores its important role in many fundamental genomic processes. While, as suggested here, the precise molecular components generating this structure may be different, the underlying basic structural features appear to be well conserved. Future work designed to characterize the physical and dynamic details of these basic features may eventually lead to an understanding of the underlying mechanisms in the various genomic processes that are conserved from insects to mammals.

## Materials and Methods

### Cell culture and synchronization

Drosophila late embryonic S2R+ cells were grown in Schneider's medium (Invitrogen) with 10% heat-inactivated fetal bovine serum (FBS) (BI) at 25 °C. Cells were synchronized at G1/S by incubating with 1 μM Hydroxyurea (HU) for 18 h (23).

### Hi-C library preparation and data processing

Hi-C libraries of two biological replicates for both asynchronous and G1/S arrested cells were generated utilizing the *in situ* Hi-C method (24) with minor modifications. Briefly, nuclei released from 10 million crosslinked cells were digested with DpnII (NEB). After end-repair and ligation, the biotin-labeled chimeric molecules were fragmented with Cavrios M220, and the fragments between 300 bp to 500 bp were selected for the generation of library. The libraries were prepared using NEBNext Ultra DNA library prep kit (#E7370, NEB) according to the manufacturer’s instructions with minor modifications (Supplementary Materials). The libraries were then sequenced using the Illumina X10 platform.

The Hi-C reads were iteratively mapped to the dm3 *Drosophila melanogaster* reference genome using bowtie2 (v2.2.9) (Supplementary Materials). After filtering, the valid contact matrix was normalized using ICE as described (40). After normalization, domains were annotated using the software Armatus (25) with the scaling parameter, gamma, set to 0.9. However, visual inspection of the analyzed data revealed that some domain borders were immediately adjacent to regions with no reads, suggesting that these locations may have been defined as borders owing to absence of reads in the adjacent region. Thus, we scanned through the analyzed data and, where there was a domain boundary adjacent to a read with no reads, we re-analyzed the data using a gamma value of 0.6.

For the calculation of the number of TADs within the super-TADs and inter-super-TADs, we established a threshold value of 75% of the domain length for inclusion in either a super-TAD or inter-super-TAD.

#### Calculation of the extent of enrichment of insulators at TAD borders

All peaks and normalized signal tracks for each insulator protein were obtained from the modENCODE database. For each domain boundary, we first identified the boundary center as the midpoint between the end position of the upstream domain and the start position of the downstream domain. For each insulator protein, we calculated the average occupancy value within 4 kb of each boundary center using an 80 bp window. We used the same method to calculate the background values by randomly changing the positons of the border centers over the entire genome. The ratios of values obtained from actual boundary centers to that from randomly shuffled centers were used to evaluate the enrichment of insulator proteins at the boundaries.

For the evaluation of the co-localization of insulator proteins with TAD borders, we considered any insulator protein peak localized within 2 kb of the domain boundary as co-localized with that boundary.

#### Prediction of domain boundaries

We used the function linear_model.LogisticRegression from the Python package scikit-learn (v0.18) to implement a logistic regression model similar to that described (19) to predict the domain boundaries using different combinations of epigenetic and insulator markers. The income variables were Z-transformed signals of different markers for each fragment from the modENCODE database, with an output value of 0 indicating an intra-domain fragment and a value of 1 indicating a border-related fragment. Training sets and test sets were separated randomly with equal sizes using the cross_validation.train_test_split function. The Receiver Operating Characteristic curves (ROC curves) and Area Under Curve (AUC) values were calculated using the functions metrics.roc_curve and metrics.auc from scikit-learn.

#### Determination of the DNA condensation within TADs

We calculated the average contact frequency of all pairs of fragments located inside each TAD as a measure of DNA condensation. The restriction fragments with no ligation products were removed from this calculation. To avoid complications arising from comparing domains of significantly different sizes, we compared only those domains whose size is between 5 kb to 20 kb, since for this range, there are a sufficient number of domains of each type.

#### Calculation of the enrichment of TAD-TAD interactions

For the evaluation of TAD-TAD interactions, an enrichment ratio matrix was first calculated by dividing the contact number of each pair of fragments by the average contact number of all pairs of fragments that have the same interaction distance, binning the distances using a 200 bp window. An average enrichment ratio was then calculated for each pair of TADs by averaging all the enrichment ratios of all pairs of fragments localized in this pair of TADs.

#### Data analysis of Hi-C data of human GM12878 cells

We downloaded the GM12878 Hi-C data from the GEO database with accession number GSE63525. We determined the normalized Hi-C heatmap using the KR normalization factors. We used the Armatus software to annotate TADs in the 1 kb resolution data using a gamma value of 0.7. We also annotated large TADs at 5 kb resolution using different gammas (0.6 - 1.0) and the majority (70.9%) of boundaries of the TADs identified by Rao el at (24) were located within 2 bins of the Armatus domain boundaries.

## Acknowledgements

We thank Dr. Hua Li and Chuansheng Hu for useful discussions. This work was supported by grants from the National Natural Science Foundation of China (Nos. 11374207, 31370750, 21273148, 31670722, 81627801, and 21303104) and Shanghai Jiao Tong University.

## References

Dixon JR, et al. (2015) Chromatin architecture reorganization during stem cell differentiation. Nature 518(7539):331–336.

Li G & Zhu P (2015) Structure and organization of chromatin fiber in the nucleus. FEBS letters 589(20 Pt A):2893–2904.

Pombo A & Dillon N (2015) Three-dimensional genome architecture: players and mechanisms. Nature reviews. Molecular cell biology 16(4):245–257.

Pope BD, et al. (2014) Topologically associating domains are stable units of replication-timing regulation. Nature 515(7527):402–405.

Wei Z, et al. (2013) Biological implications and regulatory mechanisms of long-range chromosomal interactions. The Journal of biological chemistry 288(31):22369–22377.

Wu J, et al. (2016) The landscape of accessible chromatin in mammalian preimplantation embryos. Nature 534(7609):652–657.

Dekker J, Rippe K, Dekker M, & Kleckner N (2002) Capturing chromosome conformation. Science 295(5558):1306–1311.

Dostie J, et al. (2006) Chromosome Conformation Capture Carbon Copy (5C): a massively parallel solution for mapping interactions between genomic elements. Genome research 16(10):1299–1309.

Lieberman-Aiden E, et al. (2009) Comprehensive mapping of long-range interactions reveals folding principles of the human genome. Science 326(5950):289–293.

Dixon JR, et al. (2012) Topological domains in mammalian genomes identified by analysis of chromatin interactions. Nature 485(7398):376–380.

Nora EP, et al. (2012) Spatial partitioning of the regulatory landscape of the X-inactivation centre. Nature 485(7398):381–385.

Bonev B & Cavalli G (2016) Organization and function of the 3D genome. Nature reviews. Genetics 17(11):661–678.

Busslinger GA, et al. (2017) Cohesin is positioned in mammalian genomes by transcription, CTCF and Wapl. Nature 544(7651):503–507.

Zuin J, et al. (2014) Cohesin and CTCF differentially affect chromatin architecture and gene expression in human cells. Proceedings of the National Academy of Sciences of the United States of America 111(3):996–1001.

Narendra V, et al. (2015) CTCF establishes discrete functional chromatin domains at the Hox clusters during differentiation. Science 347(6225):1017–1021.

Seitan VC, et al. (2013) Cohesin-based chromatin interactions enable regulated gene expression within preexisting architectural compartments. Genome research 23(12):2066–2077.

Sofueva S, et al. (2013) Cohesin-mediated interactions organize chromosomal domain architecture. The EMBO journal 32(24):3119–3129.

Cubenas-Potts C, et al. (2017) Different enhancer classes in Drosophila bind distinct architectural proteins and mediate unique chromatin interactions and 3D architecture. Nucleic acids research 45(4):1714–1730.

Ulianov SV, et al. (2016) Active chromatin and transcription play a key role in chromosome partitioning into topologically associating domains. Genome research 26(1):70–84.

Pandey UB & Nichols CD (2011) Human disease models in Drosophila melanogaster and the role of the fly in therapeutic drug discovery. Pharmacological reviews 63(2):411–436.

Naumova N, et al.(2013) Organization of the mitotic chromosome. Science 342(6161):948-953.

Schneider I (1972) Cell lines derived from late embryonic stages of Drosophila melanogaster. Journal of embryology and experimental morphology 27(2):353–365.

Graham AC, Kiss DL, & Andrulis ED (2009) Core exosome-independent roles for Rrp6 in cell cycle progression. Molecular biology of the cell 20(8):2242–2253.

Rao SS, et al. (2014) A 3D map of the human genome at kilobase resolution reveals principles of chromatin looping. Cell 159(7):1665–1680.

Filippova D, Patro R, Duggal G, & Kingsford C (2014) Identification of alternative topological domains in chromatin. Algorithms for molecular biology: AMB 9:14.

Liang J, et al. (2014) Chromatin immunoprecipitation indirect peaks highlight long-range interactions of insulator proteins and Pol II pausing. Molecular cell 53(4):672–681.

Schwartz YB, et al. (2012) Nature and function of insulator protein binding sites in the Drosophila genome. Genome research 22(11):2188–2198.

Negre N, et al. (2010) A comprehensive map of insulator elements for the Drosophila genome. PLoS genetics 6(1):e1000814.

Phillips-Cremins JE, et al. (2013) Architectural protein subclasses shape 3D organization of genomes during lineage commitment. Cell 153(6):1281–1295.

Vogelmann J, et al. (2014) Chromatin insulator factors involved in long-range DNA interactions and their role in the folding of the Drosophila genome. PLoS genetics 10(8):e1004544.

Boettiger AN, et al. (2016) Super-resolution imaging reveals distinct chromatin folding for different epigenetic states. Nature 529(7586):418–422.

Kharchenko PV, et al. (2011) Comprehensive analysis of the chromatin landscape in Drosophila melanogaster. Nature 471(7339):480–485.

Hou C, Li L, Qin ZS, & Corces VG (2012) Gene density, transcription, and insulators contribute to the partition of the Drosophila genome into physical domains. Molecular cell 48(3):471–484.

Sexton T, et al. (2012) Three-dimensional folding and functional organization principles of the Drosophila genome. Cell 148(3):458–472.

Dixon JR, Gorkin DU, & Ren B (2016) Chromatin Domains: The Unit of Chromosome Organization. Molecular cell 62(5):668–680.

Bailey SD, et al. (2015) ZNF143 provides sequence specificity to secure chromatin interactions at gene promoters. Nature communications 2:6186.

Van Bortle K, et al. (2014) Insulator function and topological domain border strength scale with architectural protein occupancy. Genome biology 15(6):R82.

Gorkin DU, Leung D, & Ren B (2014) The 3D genome in transcriptional regulation and pluripotency. Cell stem cell 14(6):762–775.

Li H, et al. (2016) Ultra-deep sequencing of ribosome-associated poly-adenylated RNA in early Drosophila embryos reveals hundreds of conserved translated sORFs. DNA research : an international journal for rapid publication of reports on genes and genomes 23(6):571–580.

Imakaev M, et al. (2012) Iterative correction of Hi-C data reveals hallmarks of chromosome organization. Nature methods 9(10):999–1003.

